# Tfam knockdown results in reduction of mtDNA copy number, OXPHOS deficiency and abnormalities in zebrafish embryos

**DOI:** 10.1101/843318

**Authors:** Auke BC Otten, Rick Kamps, Patrick Lindsey, Mike Gerards, Hélène Pendeville-Samain, Marc Muller, Florence HJ van Tienen, Hubert JM Smeets

## Abstract

High mitochondrial DNA (mtDNA) copy numbers are essential for oogenesis and embryogenesis and correlate with fertility of oocytes and viability of embryos. To understand the pathology and mechanisms associated with low mtDNA copy numbers, we knocked down mitochondrial transcription factor A (*Tfam*), a regulator of mtDNA replication, during early zebrafish development. Reduction of *Tfam* using a splice-modifying morpholino (MO) resulted in a 42%±4% decrease in mtDNA copy number in embryos at 4 days post fertilization. Morphant embryos displayed abnormal development of the eye, brain, heart and muscle, as well as a 50%±11% decrease in ATP production. Transcriptome analysis revealed a decrease in protein-encoding transcripts from the heavy strand of the mtDNA. In addition, various RNA translation pathways were increased, indicating an upregulation of nuclear and mitochondria-related translation. The developmental defects observed were supported by a decreased expression of pathways related to eye development and haematopoiesis. The increase in mRNA translation might serve as a compensation mechanism, but appears insufficient during prolonged periods of mtDNA depletion, highlighting the importance of high mtDNA copy numbers for early development in zebrafish.

**SUMMARY STATEMENT:** The first tuneable zebrafish model used to characterize the effect of a reduced mtDNA copy number and resulting OXPHOS deficiency on zebrafish embryonic development.

## INTRODUCTION

Mitochondria are responsible for producing the majority of cellular energy in the form of ATP. Depending on the tissue type and energy requirement, a cell can contain up to thousands of mitochondria, each having multiple copies of mitochondrial DNA (mtDNA) (Smeets 2013). Of all cell-types, mtDNA copy numbers are the highest in oocytes, ranging between 100,000 and 400,000 copies in mammals, such as rodents, cows and humans, and up to 16,5 million copies in zebrafish (Otten and Smeets 2015). The high mtDNA copy number in oocytes is established by an initial reduction during embryogenesis, called the mitochondrial bottleneck, followed by clonal expansion of the mtDNA during oogenesis and appears to be important for successful fertilization and embryogenesis (Santos et al. 2006). In mice, the oocyte mtDNA copy number should be sufficient for normal development until implantation at day 4, and it has been demonstrated that oocytes with less than 50,000 mtDNA copies fail to resume development after implantation (Ebert et al. 1988; Wai et al. 2010). This negative correlation between mtDNA copy number and developmental competence of embryos has also been suggested for human oocytes (Yamamoto et al. 2010).

Being a key activator of mtDNA replication and transcription, mitochondrial transcription factor A (TFAM), a protein of the high mobility group box-family (Parisi et al. 1993), is crucial in the regulation of mtDNA copy number (Larsson et al. 1998). A direct relation between the mtDNA copy number and the total TFAM protein level has been demonstrated in mice embryos with a heterozygous or homozygous disruption of the *Tfam* gene (Ekstrand et al. 2004). Homozygous TFAM knockout (KO) mouse embryos displayed a strong mtDNA depletion with severely reduced functioning of the electron transport chain (ETC) and died after gastrulation and implantation, while heterozygous KO TFAM mice had moderately reduced mtDNA levels and respiratory chain deficiency, which was strongest in the developing heart (Larsson et al. 1998). Together, these studies demonstrate the importance of a sufficient mtDNA copy number during oogenesis and embryogenesis, but the mechanism by which a reduced mtDNA copy number affects embryogenesis is currently unknown.

Studying embryonic development in zebrafish overcomes ethical and practical obstacles associated with the use of human or mouse embryos. Zebrafish are vertebrates, >70% of human genes have at least one zebrafish orthologue, and the major tissues and organs are the same. Also, zebrafish are transparent during early development and they have a high number of offspring. Organs develop within 5 days post fertilization and gene knockdown during early embryogenesis can be performed by injection of morpholino antisense oligonucleotides (MOs) (Pauli et al. 2015). Previously, we showed that the mitochondrial bottleneck during early development is highly conserved between zebrafish and mammals (Otten et al. 2016). In addition, the zebrafish *Tfam* protein is functionally homologous to its human counterpart and is expressed ubiquitously from the earliest stages of development (Artuso et al. 2012). In this study we performed Tfam knockdown during zebrafish embryogenesis in order to reduce the mtDNA copy number and assess its effect on OXPHOS function and embryonic development and to determine the underlying mechanisms.

## RESULTS

### Tfam knockdown results in decreased mtDNA content and mitochondrial ATP production

To establish decreased mtDNA content, we microinjected an antisense splice-MO targeting *tfam*-mRNA in zebrafish embryos. Using RT-PCR and Sanger sequencing of cDNA from *tfam* MO-injected 4 dpf embryos, we showed that the *tfam* splice-MO causes a 128 base-pair deletion of exon 2 (c.84_211del), predicted to cause a frameshift and a premature stop codon (p.(Cys29Hisfs*36)) (Fig. 1A, Supplementary Fig. S1A). Quantitative PCR analysis of *tfam* exon 5 and exon 6-7 showed respectively 59%±1% and 60%±2% decrease in expression in *tfam* splice-MO injected embryos at day 4 (n=8/group). qPCR analysis of *tfam* exon 2 expression showed a reduction of 80±1% at day 4 in zebrafish injected with 2 ng *tfam* splice-MO compared to the Ctrl-MO (Supplementary Fig. S1B). This indicates that ~60% of *tfam* RNA is subjected to nonsense-mediated decay and that of the residual ~40% *tfam* RNA, only half is exon 2-containing wild-type RNA (20% of total tfam/tbp). Next, we assessed the mtDNA copy number in non-injected, Ctrl-MO injected and *tfam* splice-MO injected embryos (n=10 in each group). At 4 dpf, a significant, 42%±4% (SEM) decrease in the mtDNA copy number was observed in embryos injected with either 2 ng or 4 ng *tfam* splice-MO (Fig. 1B). In contrast, no significant differences in mtDNA copy number were apparent between groups at 2 dpf (Supplementary Fig. S1C). To assess if decreased mtDNA copy number also affected mitochondrial capacity, we measured the oxygen consumption rate (OCR) at 4 dpf. As shown in Fig. 1C, both the OCR of the basal respiration and the ATP production were significantly decreased in the 2ng *tfam* splice-MO treated zebrafish (n=10) compared with the 2ng control-MO treated group (n=9) (p<0.05) by respectively 40%±8% and 50%±11% (SEM).

**Fig. 1.**
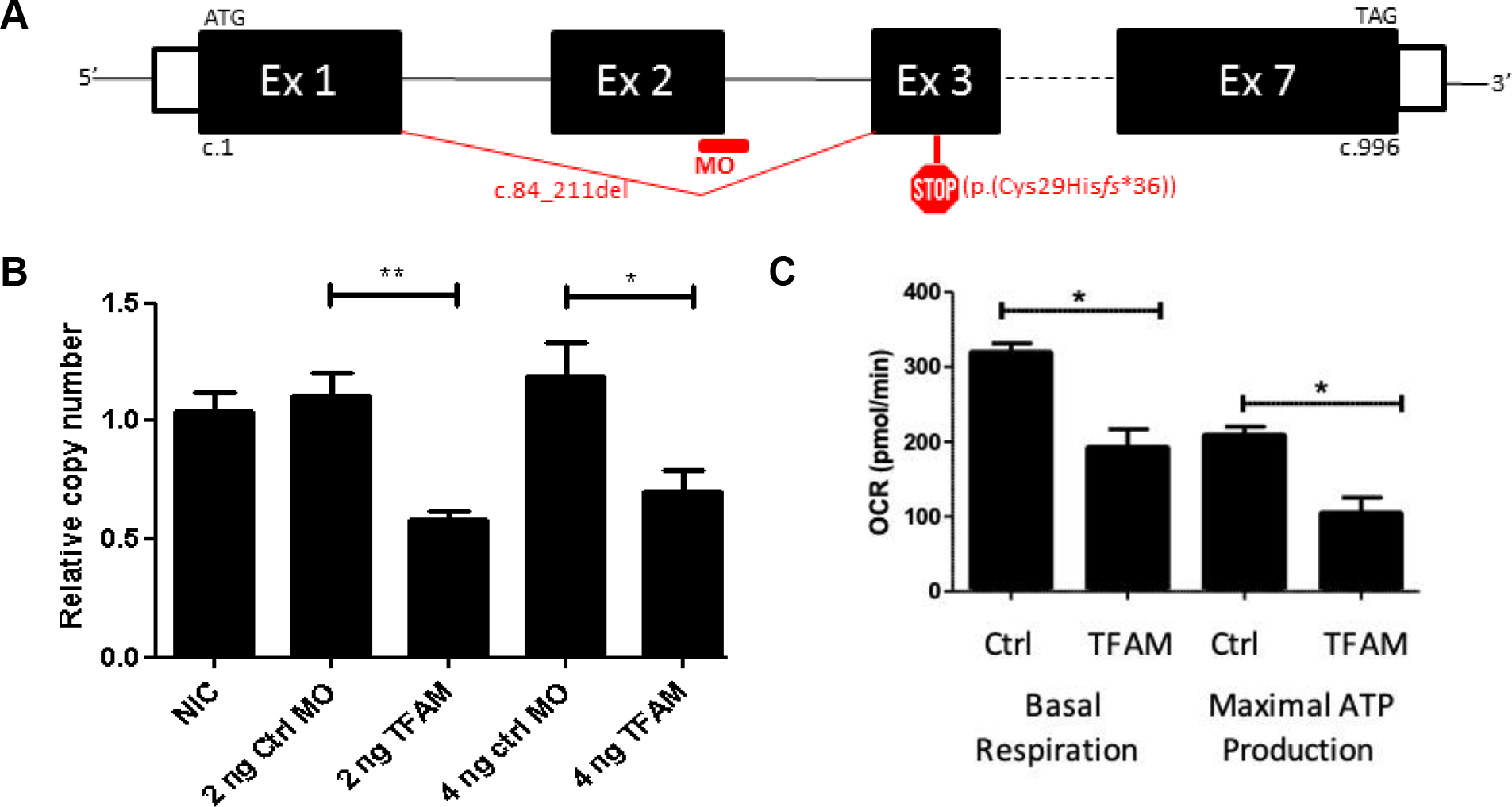
Knockdown of TFAM in zebrafish embryos. Zebrafish embryos injected with indicated amount of either Control-morpholino (Ctrl-MO) or *tfam* splice-morpholino (TFAM-MO) at 1 hpf and analysed at 4 dpf. (A) *tfam* splice-morpholino (MO) at the 3’ splice site of exon 2 causes deletion of exon 2, which predicts a frame-shift and premature stop codon (p.(Cys29Hisfs^*^36)) (figure is not on scale). (B) The relative copy number assessed by mitochondrial ND1 / nuclear B2M ratio. Data are normalized to the NIC embryos. Bars indicate mean values with SEM. *P*-values are calculated using *Bonferroni’s Multiple Comparison* Test to assess copy number, * *P*-value < 0.05, *002A; *P*-value < 0.01. (C) The oxygen consumption rate (OCR) measured by Seahorse XF24 in order to assess basal respiration and maximal ATP production capacity at 4 dpf using 2ng MO. The statistical analysis was performed by using a student T-test *(P* < 0.05).

### Decreased mtDNA contents cause brain, eye, heart, and muscle abnormalities

At 4 dpf, morphant embryos displayed morphological abnormalities compared to control embryos. The macroscopic phenotype included overall oedema, curved tails, necrotized yolk sacs and small eyes (Supplementary Fig. S1D). Fish injected with 4 ng *tfam* splice-MO were more severely affected, as they had a higher count for oedema, curved tails, necrosed yolk sac, small eyes, and developmental delay. The percentage of dead embryos was <1% for both concentrations of *tfam MO*-injections at 4 dpf (Supplementary Fig. S1D). Strikingly, all observed pathologies manifested at 3 or 4 dpf, whereas development was apparently normal for *tfam* splice-MO injected embryos at 2 dpf upon macroscopic inspection. These morphological changes parallel the decrease in mtDNA copy number that is observed at 4 dpf (Fig. 1B), but not at 2 dpf (Supplementary Fig. S1C). Since the decrease in mtDNA content was comparable with 4 ng *tfam* splice-MO, the dosage of 2 ng *tfam* splice-MO was used in all following experiments, reducing the risk of non-specific observations.

Microscopic evaluation of zebrafish depleted of mtDNA showed that they were less well-developed than control embryos, as demonstrated by a decreased brain size with no or less well-developed cerebellum (Fig. 2B). In addition, morphant embryos had smaller eyes and of which the different layers were less-well organized compared to controls (Fig. 2D). Also, the organization of the myotomic area was less compact with the skeletal muscle fibres being thinner and disorganized (Fig. 2F). Finally, morphant fish displayed marked pericardial oedema, alongside with a dilated non-looped heart (Fig. 2H).

**Fig. 2.**
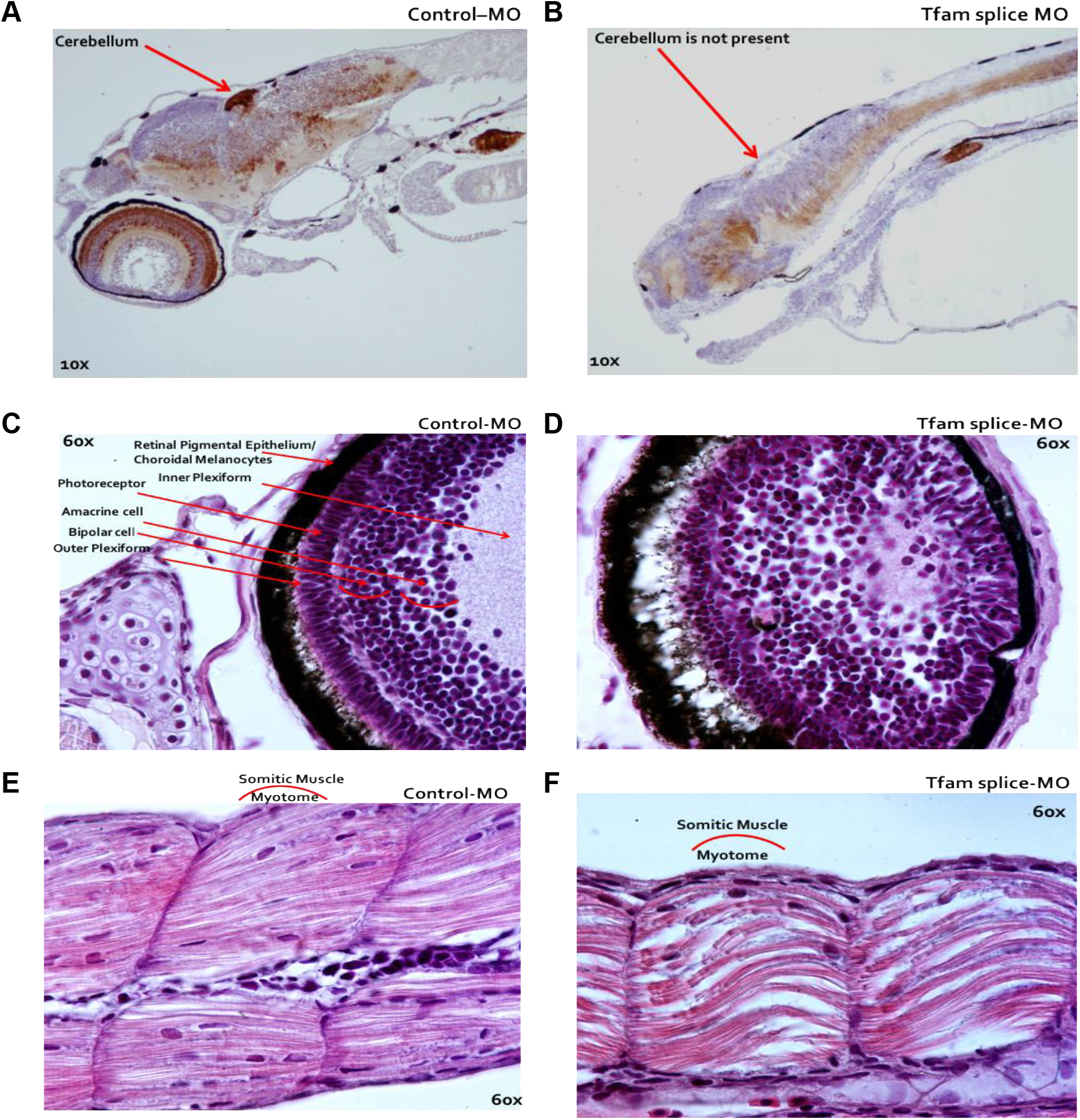
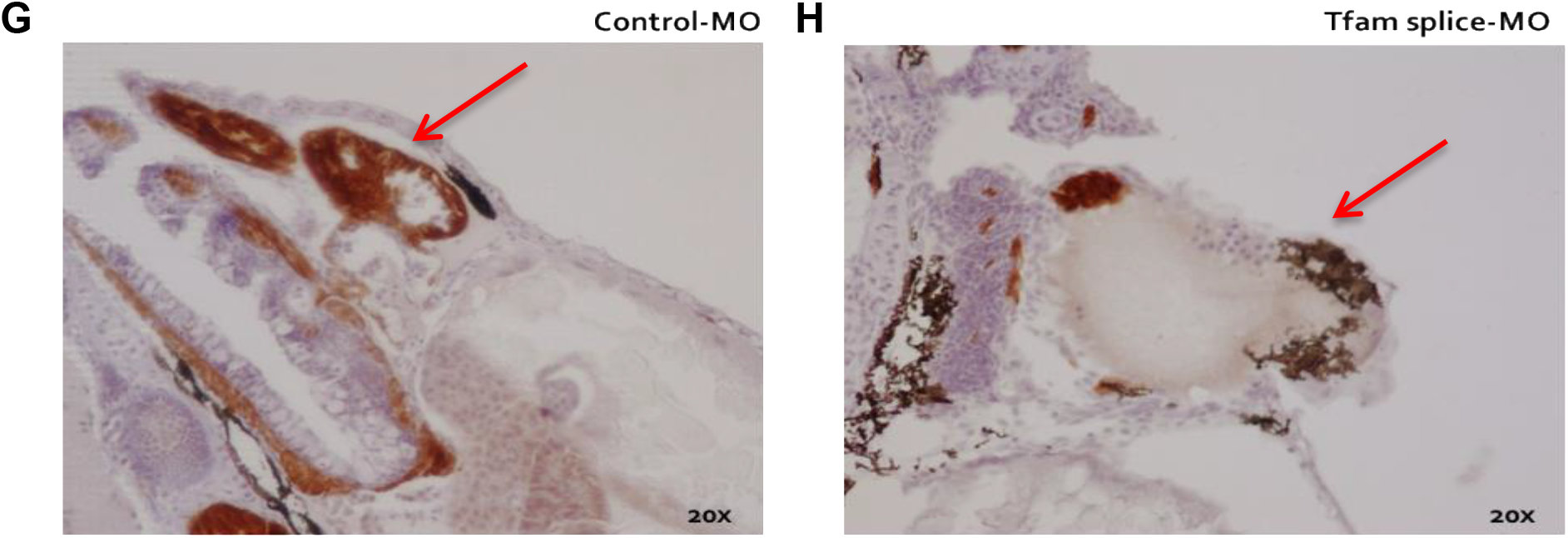
Microscopy of zebrafish embryos following TFAM knockdown. Image (A-H): Microscopic analysis following serial HE-staining of zebrafish embryos at 4 dpf, which were injected at 1 hpf with either 2 ng *Tfam* splice-MO or 2 ng Ctrl-MO. (B) The cerebellum was missing in the phenotype or a delayed development of this cerebellum in these fish.(D) The eyes were smaller in the phenotype and the eye layers were less developed. (F) A muscle phenotype was clearly visible in the dorsal fin, especially in the myotomic area, which was less compact in these somatic muscles. (H) The fish displayed a marked pericardial oedema (red arrow). They did not develop a normally looped heart as in Ctrl-MO (2G), but the hearts were dilated in the phenotypes (2H).

### Genes and pathways altered by Tfam knockdown

To characterize the underlying pathogenic processes due to *tfam* knockdown and subsequent decrease in mtDNA copy number, we compared global gene expression profiles at 4 dpf of zebrafish injected with 2 ng of either *tfam*-MO (n=12) or Ctrl-MO (n=12). A total of 19,459 transcripts were present on the array (Supplementary Table S2), of which 16,631 (85.5%) had a signal intensity higher than twice the background signal in both conditions or a signal that was at least three times higher than the background signal in one condition. Of these, 3,158 transcripts (19.3%) were differentially expressed between the two groups with a fold change (FC) >50%. Expression of 2,063 transcripts (64.3%) was increased (FC ≥1.50), while expression of 1,145 transcripts (35.7%) was decreased in the *tfam* splice-MO injected embryos (FC ≤0.67)). In order to identify significantly altered pathways by using gene ontology analysis, Protein ANalysis THrough Evolutionary Relationships (PANTHER) was used. PANTHER analysis included 17,395 transcripts. A total of 18 significantly enriched GO biological processes (FDR <0.05) were identified and these were manually grouped into four groups, namely: RNA processing, ribosome biogenesis, drug metabolism, and visual perception (Table 1, and Supplementary Table S4). We also listed the mitochondria-related processes, characterized by mitochondria-related GO terms (Table 2, Supplementary Table S5). Although these processes were not significantly altered, the expression of the 8 mtDNA protein encoding genes was significantly decreased, all encoded by the H-strand, whereas the one transcript on the L-strand was unaltered. Four transcripts were not present (atp6, atp8, cox 3, and cytb) in the analysis (Supplementary Table S3). Notably, we observed that the GO terms analysis did not contain all mitochondrial genes known to be involved in these processes as expected, which obviously limits the analysis of mitochondria-related processes.

**Table 1.** Significantly altered GO biological process terms

| Gene set | Description | O-T | C-T | Fold Enrichment | FDR-T |
| --- | --- | --- | --- | --- | --- |
| **RNA processing** |  |  |  |  |  |
| GO:0034475 | U4 snRNA 3'-end processing | 8 | 8(Cm=0,Cn=8) | 6.09 | 3.2E-02 |
| GO:0034470 | ncRNA processing | 197 | 72(Cm=14,Cn=58) | 2.22 | 4.3E-06 |
| GO:0034660 | ncRNA metabolic process | 260 | 88(Cm=30,Cn=58) | 2.06 | 5.2E-06 |
| GO:0006396 | RNA processing | 426 | 106(Cm=15,Cn=91) | 1.52 | 2.4E-02 |
| GO:0016072 | rRNA metabolic process | 117 | 47(Cm=6,Cn=41) | 2.44 | 1.1E-04 |
| GO:0008033 | tRNA processing | 73 | 27(Cm=7,Cn=20) | 2.25 | 3.8E-02 |
| GO:0006399 | tRNA metabolic process | 118 | 44(Cm=22,Cn=22) | 2.27 | 1.4E-03 |
| GO:0009451 | RNA modification | 88 | 32(Cm=9,Cn=23) | 2.21 | 2.6E-04 |
| **Ribosome biogenesis** |  |  |  |  |  |
| GO:0006364 | rRNA processing | 105 | 43(Cm=6,Cn=37) | 2.49 | 2.0E-04 |
| GO:0042254 | Ribosome biogenesis | 162 | 60(Cm=14,Cn=46) | 2.25 | 4.3E-05 |
| GO:0022613 | Ribonucleoprotein complex biogenesis | 239 | 69(Cm=14,Cn=55) | 1.76 | 1.0E-02 |
| **Drug metabolism** |  |  |  |  |  |
| GO:0042737 | Drug catabolic process | 85 | 33(Cm=2,Cn=31) | 2.36 | 1.1E-02 |
| GO:0071028 | Nuclear mRNA surveillance | 8 | 8(Cm=0,Cn=8) | 6.09 | 3.6E-02 |
| GO:0071027 | Nuclear RNA surveillance | 8 | 8(Cm=0,Cn=8) | 6.09 | 3.4E-02 |
| GO:0071025 | RNA surveillance | 9 | 9(Cm=0,Cn=9) | 6.09 | 2.3E-02 |
| GO:0016075 | rRNA catabolic process | 11 | 9(Cm=0,Cn=9) | 4.98 | 3.6E-02 |
| **Visual perception** |  |  |  |  |  |
| GO:0007601 | Visual perception | 83 | 31(Cm=2,Cn=29) | 2.27 | 2.3E-02 |
| GO:0050953 | Sensory perception of light stimulus | 88 | 31(Cm=2,Cn=29) | 2.14 | 3.5E-02 |

Total analysis (T): 2,856 significantly changed unique Entrez Gene IDs and 17,395 unique array reference Entrez Gene IDs were used to identify overrepresented GO terms biological process. GO terms with $FDR < 0.05$ were considered to be significantly changed. Displayed are counted number of genes (C-T= Cmitochondrial (Cm),Cnuclear (Cn)), Over-represented genes (O-T), Enriched number of genes (Fold Enrichment), Binomial test is showing False Discovery Rate (FDR).

**Table 2.** Mitochondria-related GO biological processes

| Gene set | Description | O-T | C-T | Fold Enrichment | FDR-T |
| --- | --- | --- | --- | --- | --- |
| **Mitochondrial processes** |  |  |  |  |  |
| GO:0007005 | mitochondrion organization | 152 | 15 | 0.60 | 6.7E-01 |
| GO:0007007 | inner mitochondrial membrane organization | 20 | 2 | 0.61 | 9.7E-01 |
| GO:0007008 | outer mitochondrial membrane organization | 1 | 0 |  | 9.9E-01 |
| GO:0006390 | mitochondrial transcription | 8 | 2 | 1.52 | 9.5E-01 |
| GO:0032543 | mitochondrial translation | 27 | 8 | 1.80 | 1.0E00 |
| GO:0006626 | protein targeting to mitochondrion | 32 | 1 | 0.19 | 7.6E-01 |
| GO:0070585 | protein localization to mitochondrion | 32 | 1 | 0.19 | 7.6E-01 |
| GO:0042775 | mitochondrial ATP synthesis coupled electron transport | 41 | 7 | 1.04 | 5.1E-01 |
| GO:0006119 | oxidative phosphorylation | 46 | 9 | 1.19 | 3.5E-01 |
| GO:0033108 | mitochondrial respiratory chain complex assembly | 25 | 2 | 0.49 | 9.6E-01 |
| GO:0015986 | ATP synthesis coupled proton transport | 20 | 1 | 0.30 | 1.6E-01 |
| GO:0006979 | response to oxidative stress | 70 | 9 | 0.78 | 2.9E-01 |
| GO:0006839 | mitochondrial transport | 73 | 6 | 0.50 | 8.5E-01 |
| GO:1990542 | mitochondrial transmembrane transport | 44 | 3 | 0.41 | 10.0E-01 |
| GO:0008053 | mitochondrial fusion | 12 | 3 | 1.52 | 9.4E-01 |
| GO:0000266 | mitochondrial fission | 16 | 1 | 0.38 | 1.0E00 |
| GO:0000422 | autophagy of mitochondrion | 27 | 2 | 0.45 | 9.0E-01 |
| GO:0000002 | mitochondrial genome maintenance | 6 | 1 | 1.01 | 8.9E-01 |
| GO:0009117 | nucleotide metabolic process (DNA replication) | 282 | 46 | 0.99 | 5.2E-01 |

Total analysis (T): 2,856 significantly changed unique Entrez Gene IDs and 17,395 unique
array reference Entrez Gene IDs were used to identify overrepresented GO terms biological process. GO terms with FDR > 0.05 were considered to be unaltered mitochondrial process. Displayed were counted number of genes (C-T), Over-represented genes (O-T), Enriched number of genes (Fold Enrichment), Binomial test is showing False Discovery Rate (FDR).

## DISCUSSION

A high mtDNA copy number is critical for normal embryonic development. A key regulator of the mtDNA copy number is TFAM, as TFAM initiates mtDNA replication by its capability to wrap, bend and unwind the mtDNA (Ekstrand et al. 2004), a function conserved in zebrafish (Howe et al. 2013). In order to study the effect of a decreased mtDNA copy number during zebrafish embryonic development, we performed *tfam* knock-down using a splice-morpholino (MO). We demonstrated that the *tfam* splice-MO induces skipping of exon 2, predicting a frameshift and a premature stop codon on protein level (p.(Cys29Hisfs*36)) (Fig. 1A), and that wild-type *tfam* RNA expression is reduced to 20% of control values. At 3 and 4 dpf, morphant zebrafish have a decreased mtDNA copy number and are less well developed. This correlates with the previous observations that in wild-type zebrafish larvae an increase in mtDNA copy number only becomes apparent between 2 and 3 dpf (Otten et al. 2016) indicating that, in the absence of mtDNA replication, mtDNA copy number becomes critically low at 3-4 dpf and an increase is required for embryonic tissue differentiation and organogenesis.

OXPHOS capacity was significantly lower in 4 dpf morphant embryos. This occurs at a critical phase of development, when there is a high energy demand and a metabolic switch from glycolysis to OXPHOS takes place, essential for healthy organogenesis (Hance et al. 2005; Stackley et al. 2011). This necessity is illustrated by the pericardial oedema and reduced brain size we observed in morphant zebrafish (Fig. 2B, 2H). The heart and the brain are the first organs to develop (Glickman and Yelon 2002) and therefore the first to rely on OXPHOS, making them more prone to OXPHOS defects. Moreover, as in a normal heart >30% of the cardiac muscle volume is occupied by mitochondria, the deficit of mtDNA (and subsequently mitochondria) also hampers proper alignment of the muscle fibres in the developing heart, creating structural problems as well (Ventura-Clapier et al. 2011). In mice, a *Tfam* KO model has been developed, allowing comparison of our zebrafish data with homozygous and heterozygous KO mice. A complete failure of organogenesis and embryonic death has been observed in homozygous *Tfam* KO mice with abolished ETC function, due to a severe mtDNA depletion (Larsson et al. 1998). Mice with 50% *Tfam* expression were viable, but showed a respiratory chain deficiency in the developing heart and a dilated cardiomyopathy and conduction block later in life (Larsson et al. 1998; Powell et al. 2015). The severity of our model seems to fit somewhere in between, as wild-type *tfam* levels are around 20% of control level at the day of analysis. Noticeably, MOs establish only a transient knockdown, which is not the case for a homozygous or heterozygous knockout of the gene, prohibiting a proper comparison later in life.

In order to unravel the biological processes that were altered due to decreased *tfam* expression which resulted in a decrease in mtDNA copy number, we performed global gene expression analysis. The processes showing the largest gene expression differences could be categorised into four groups, being RNA processing, ribosome biogenesis, drug metabolism, and visual perception (Table 1). The first two groups show that RNA translation in general is upregulated. Processes in the group drug metabolism, like nuclear (m)RNA surveillance, nuclear RNA surveillance and rRNA catabolic process, also reflect an upregulation at the translation level. From the gene list (SupplementaryTable S4), it can be observed that the translation of both nuclear and mitochondrial transcripts is increased, the latter including genes encoding for mitochondrial ribosomal proteins, mitochondrial aminoacyl-tRNA synthetases (Sylvester et al. 2004; Konovalova and Tyynismaa 2013), tRNA processing and modification enzymes, translation activators and mRNA stability factors and translational initiation, elongation and termination factors (Boczonadi and Horvath 2014). In line with this, also 8 out of 9 exosome components were significantly upregulated in *tfam* splice-MO zebrafish. Biological processes comprising mitochondrial respiratory chain components were not significantly changed, except that the expression of 8 out of the 9 mtDNA protein encoding transcripts was decreased, all encoded by the H-strand, whereas the one transcript on the L-strand was unaltered. Most likely this reflects a switch or preferable transcription from the Light-Strand Promoter (LSP), which is induced by low amounts of *tfam* and which also primes H-strand replication (Shutt et al. 2011), but which in our case fails to increase mtDNA copy number due to the persisting lack of TFAM. Also, within the biological process drug catabolism, several downregulated genes are linked to mitochondrial function, including haemoglobin production. As haem is produced by mitochondria (Wagener et al. 2003), this could be explained by the decreased amount of mtDNA and subsequently mitochondria at the onset of definitive haematopoiesis, initiating between 1 to 2 dpf (Murayama et al. 2006; Bertrand et al. 2007; Zhang and Rodaway 2007; Zhang and Hamza 2018). Obviously, deficiencies in haematopoiesis during development might have downstream effects as well, comparable to the OXPHOS deficiency. Other mitochondria-related genes, such as OXCT1a, OXCT1b, members of CYP450 family 2 and GPX1a were also downregulated in *tfam* splice-MO-injected zebrafish. These genes are related to metabolism of chemicals and other metabolites, suggesting a build-up of toxins and a reduction in signalling molecules.

It is striking that the general reaction of the organism to a decreased mtDNA copy number is an overall increase in RNA translation and translation regulation processes. Increasing the efficiency of the translation of available mRNA could be a reaction to the lack of mtDNA and OXPHOS capacity. Also upregulation of the exosome components seems to be such a compensatory mechanism, as exosomes contain and shuttle mRNA and are important in intercellular communication and developmental patterning (Wan et al. 2012; Boczonadi et al. 2014; McGough and Vincent 2016). Future assessment of the exosome composition is needed to provide further insight into the mechanism by which decreased mtDNA copy number affects exosome signalling and composition. A reduction in mtDNA severely impacts organogenesis (Fig. 2), but is in the gene expression data at an organ or tissue level (Table 1) only apparent for visual perception, which consists of 23 eye genes that are all relevant for early development of the retina (Gestri et al. 2012). Retinal development is critical between 32 and 74 hpf (Renninger et al. 2011a; Renninger et al. 2011b), which is the timeframe when the decrease in mtDNA copy number manifests, explaining the down-regulation observed. No change in cardiac, muscle or brain nor in stress-related pathways was observed, suggesting that although pathology clearly manifested, we are still observing the processes preceding the pathology and not those resulting from it in the pathway analysis. Apparently, eye development seems to precede the development of the other organ systems or has a higher sensitivity to energy deficits. In addition, a number of temporally expressed developmental genes are downregulated in *tfam* splice-MO zebrafish at 4 dpf, such as chitinase (CHIA) 1, 2, 3 and 4. Expression of CHIA.1, CHIA.2 and CHIA.3 expression normally drastically increases at 4 and 5 dpf, while CHIA.4 is more stably expressed. Chitinases are suggested to play a role in early embryo immunity and chitinase inhibition has been shown to cause development disruption in zebrafish, characterized by hampered trunk and tail formation (Semino and Allende 2000).

In conclusion, inhibition of *tfam* expression during early zebrafish development leads to a decrease in mtDNA copy number, mtDNA encoded transcripts and a decrease OXPHOS capacity. This triggers a compensation mechanism at the level of translation initiation, but as pathology emerges, this compensation mechanism falls short. The eyes, heart and brain are the first organs of which the development is severely affected, corroborated by a decrease in haem production and haemoglobin, affecting the oxygen supply in the developing embryo.

## MATERIAL AND METHODS

### Zebrafish maintenance and procedures

Zebrafish (*Danio rerio*) were housed and raised in the zebrafish facility of the University of Liège as described before (Larbuisson et al. 2013). To retrieve eggs, wild-type adult male and female zebrafish were separated within the same breeding tank by a plastic divider the day before breeding. This separation was removed the next day after the light was turned on in order to allow natural mating and eggs were collected after spawning. Eggs were collected in Petri dishes containing 1x E3 medium for zebrafish at 28°C (580 mg/l NaCl, 27 mg/l KCl, 97 mg/l CaCl_2_·2H_2_O, 163 mg/l MgCl_2_·6H_2_O and 0.0001% methylene blue (Sigma-Aldrich), pH 7.2) (Larbuisson et al. 2013). Embryos were microscopically staged according to the embryonic development as described before (Kimmel et al. 1995). Unless stated otherwise, all reagents used in this study were obtained from Thermo Fisher Scientific.

### Tfam knockdown experiments

Antisense morpholino oligonucleotides (MO) were purchased from Gene Tools and micro-injected into one or two-cell stage embryos. A splice modifying MO was used, targeting the boundary of exon 2 and intron 2-3 of the zebrafish *tfam* gene (Ensemble: ENSDART00000092009, *tfam* splice-MO: 5’-CGGCAGATGGAAATTTACCAGGATT-3’). For controlling the effect of the MO injection, an equal concentration of a standard control-morpholino (Ctrl-MO: 5’-CCTCTTACCTCAGTTACAATTTATA-3’) was used in separate embryos of the same batch during each experiment. All MOs were dissolved in 1x Danieau buffer (58 mM NaCl, 0.7 mM KCl, 0.4 mM MgSO_4_, 0.6 mM Ca(NO_3_)_2_, and 5.0 mM HEPES pH 7.6) and 0.5% rhodamine dextran (Thermo Fisher) was added to the MO solution upon injection. At 1 hpf, 1 nl was injected into each embryo using microinjection (Bill et al. 2009). The next day, distribution of the rhodamine dextran dye was checked using fluorescence stereomicroscopy to verify proper injections. Only those embryos in which the rodamine dextran dye was visible were used for follow-up investigations.

### Quantitative and qualitative analysis of Tfam RNA

Total RNA of 2 – 4 dpf zebrafish embryos was extracted using Trizol reagent and purified using the RNeasy Mini Kit (Qiagen), according to manufacturer’s instructions. cDNA synthesis was performed with 500 ng RNA in the presence of first strand buffer, 0.75 μg oligo-dT, 0.75 μg random hexamer-primer, 50 nmol dNTPs (GE Healthcare Life Sciences), 1 U RNAse inhibitor (RNAsin, Promega) and 5 U reverse transcriptase for 60’ at 42°C, followed by 5’ at 95°C. The effect on *tfam* splicing was assessed using RT-PCR amplification of 25 ng cDNA in a PCR mix contained under standard conditions, using 0.6μM forward primer, 0.6 μM reverse primer (Supplementary Table S1). PCR conditions were 5’ denaturation at 94°C, 35 cycles of 1’at 94°C, 1’ at 58°C and 1,5’ at 72°C, followed by 7’ at 72°C. The PCR product was sequenced by conventional Sanger sequencing. *tfam* gene expression was quantified by comparing the RNA expression ratio of *Tfam* RNA to *18S* RNA. Per reaction, 2.5 ng cDNA was amplified in a 10 μl reaction containing 1x Sensimix Sybr Hi-Rox reagents (Bioline) and 375 nM of both forward and reverse primer (Supplementary Table S1). Real-time PCR was performed on an ABI7900HT using the following settings: 10’ 95°C, 40 cycles of 15” 95°C and 1’ 60°C. The statistical analysis was performed by a Student’s t-test. *p-values* <0.05 were considered significant.

### Quantification of mtDNA copy number

To determine mtDNA copy number, embryos were individually collected in sterile tubes and directly frozen without water. For DNA isolation, embryos were thawed and lysed for 4 hours at 55°C in lysis buffer containing 75 mM NaCl, 50 mM EDTA, 20 mM HEPES, 0.4% SDS and 200 μg proteinase K (Sigma) while vortexing repeatedly. DNA precipitation was performed overnight at −20°C after adding 420 μl isopropanol. Following centrifugation, the DNA pellet was washed with 70% ethanol and dissolved in Tris-EDTA buffer. The relative mtDNA abundance was quantified by measuring the steady-state amount of mitochondrial ND1 and nuclear B2M. Per reaction, 20 ng DNA was used in a 10 μl reaction containing 1x Sensimix Sybr Hi-Rox reagents (Bioline) and 375 nM of both forward and reverse primer (Supplementary Table S1). Real-time quantification was performed on an ABI7900HT as described before. Statistical analysis was carried out using one-way analysis of variance (ANOVA) followed by the Bonferroni multiple comparisons test. *P*-values <0.05 were considered significant.

### Imaging of zebrafish embryos

Zebrafish embryos were fixed at 4°C with paraformaldehyde for 4 hours and standard paraffin serial sections of 4 μm thickness were prepared for immunostaining and hematoxylin-eosin (HE)-staining. Immunohistochemistry was performed following a microwave heat-induced antigen retrieval step for four times 5’ at 650 Watt (in Tris-EDTA, pH=9.0) and was analysed with the Dako REALTM EnVisionTM Detection System, Peroxidase/DAB, Rabbit/Mouse, using a Dako automated immunostaining instrument and protocols according to the manufacturer (Agilent, Santa Clara). Antibodies and conditions used are listed in Supplementary Table S1. HE-staining was performed to reveal the underlying embryonic structures. Paraffin slides were embedded in Entellan (Merck) and protected by cover slips (Knittel Glass). Microscopic images were taken by using the Nikon Eclipse E80 Imaging System (Nikon) at different magnifications. The orientation of fish was determined according to the Zebrafish Bio-Atlas (Atlas 2019, 8 August).

### Oxygen consumption rate

At 4 dpf, chorion-free living fish with a heartbeat and active swimming behaviour upon touching were selected and collected individually. The oxygen consumption rate (OCR) was determined using the Seahorse XF24 system (Agilent/Seahorse Biosciences), according to the islet protocol of the manufacturer with 2 fish per well of a 24-well plate. Compound concentrations were adjusted to 80 μM for Oligomycin (Sigma-Aldrich) and 60 μM for Antimycin & Rotenone (Sigma-Aldrich) in the injecting assay medium (1x E3 medium). The OCR level at basal respiration (pmol/min) was calculated by the start respiration level minus antimycin/rotenone level. Maximal ATP production (pmol/min) was determined as start respiration level minus oligomycin level. The statistical analysis was performed by a Student’s t-test. *p*-values <0.05 were considered significant.

### Gene expression and pathway analysis

For gene expression studies, 12 embryos from each group were individually collected in sterile tubes. RNA from single embryos was isolated using the MagMax-96 for microarray kit (Ambion) and 200 ng RNA was labelled (Cyanine 3-CTP and Cyanine 5-CTP), fragmented and hybridized using the Low Input Quick Amp Labeling Kit, Two-colour (Ambion), according to manufacturer’s instructions. After labelling, amplified cRNA samples were purified using the RNeasy Mini kit (Qiagen) and Cyanine 3 and Cyanine 5 dye concentration, RNA absorbance ratio (260/280nm) and cRNA concentration were quantified using the nanodrop 2000C (Thermoscientific). Only samples with a yield >0.825 μg and a specific dye activity >6.0 pmol/μg were used for fragmentation and hybridization. For fragmentation, 825 ng labelled cRNA was used and the final volume was adjusted to 41.8 μl with RNAse-free water, followed by the hybridization procedure, as described by the manufacturer (Ambion). Dye-swap hybridizations (2+2) were performed on microarray slides (4×44K zebrafish V3, Agilent) using gasket slides and a hybridization chamber and incubated for 17 h at 65°C and 10 rpm in the hybridization oven (Agilent Technologies). Slides were washed with Triton X-102, freshly added to the Wash Buffers. Microarray slides were scanned using a DNA Microarray scanner with *Surescan* High-Resolution Technology (Model 2565CA, Agilent).

The arrays contained 45,220 probes. Each probe identifier was transformed to Ensembl gene IDs (ENSDARG). This resulted in 36,156 probes containing a non-empty transcript ID of which 19,459 were unique transcripts and kept for the analysis. All transcripts were analysed using a multivariate Gaussian linear regression (MVN(*μ*Σ) where *μ* is the mean, Σ is the covariance matrix 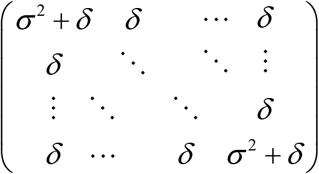, σ^2^ is the variance, and *δ* is both the extra component of variance across subjects and the common covariance among responses on the same subject) including slide differences (Slide), dye swap (Dye), background level (Bg), injection (Inj) and a random effect. The inference criterion used for comparing the models is their ability to predict the observed data, *i.e*. models are compared directly through their minimized minus log-likelihood. When the numbers of parameters in models differ, they are penalized by adding the number of estimated parameters, a form of the Akaike information criterion (AIC)(Akaike 1973). For each transcript, a model containing the relevant covariates mentioned above (E(*y*)=Slide+Dye+Bg+Inj) was fitted in order to obtain a reference AIC. Then a model containing the treatment group (Trt) was fitted (E(*y*)=Slide+Dye+Bg+Inj+Inj:Trt). The transcript under consideration was found to be differentially expressed if the AIC of this second model decreased compared to the model not containing the treatment. These statistical analysis were performed using the freely available program R(Ihaka R 1996) and the publicly available libraries ‘rmutil’ and ‘growth’(Lindsey 1999). An unbiased Gene-ontology analysis and visualization of microarray data on biological pathways was performed using PANTHER (Protein Analysis THrough Evolutionary Relationships) with the *Danio rerio* (Dr_Derby_Ensembl_80) gene product/protein database(Thomas et al. 2003),(Mi et al. 2017). Differentially expressed genes (DEGs) were mapped to unique Entrez Gene IDs and to Gene Ontology (GO) classes, using the PANTHER Overrepresentation Test (Released 2018-02-03) enrichment method of PANTHER version 13.1.

### Sample size and statistical analyses

The sample size was determined based on the mtDNA copy number, the primary outcome. It was calculated for a power of 0.8 and a statistical significance rate of 0.05. This resulted in a group size of 26 zebrafish embryos for each condition per experiment. The same groups were then used for the gene expression and morphology analysis. To minimize genetic background variability, offspring embryos were selected from the same parent zebrafish for each experiment. Except for the *tfam* splice-MO dose response analysis compared to Ctrl-MO for the mtDNA copy number where 50 zebrafish embryos were selected per condition. All collected samples were part of randomized experiments. Descriptive and group comparison analyses were carried out using Prism-GraphPad (https://www.graphpad.com/scientific-software/prism/). Student’s t-test with a significance level of 0.05 were used for all group comparisons and multiple comparison correction (Bonferroni) was applied when appropriate.

## Acknowledgements

The authors would like to acknowledge Jo M. Vanoevelen (MUMC+, Maastricht), Ellen EH Lambrichs (MUMC+, Maastricht), Sabina JV Vanherle (MUMC+, Maastricht), and Levi GK Wackers (Maastricht University, Maastricht) for their technical laboratory skills and practical contribution to this study. M.M. is a “Maître de Recherche”, supported by the “Fonds National pour la Recherche Scientifique” (FNRS). This work was further supported by the Interreg IV program of the European Council (the Alma in Silico project to M.M. and H.J.M.S.) and the European Research Area Network for Research Programmes on Rare Diseases 2 project GENOMIT (grant R 50.02.12F to M.M.). Part of this work has been made possible with the support of the Dutch Province of Limburg (M.G., and H.S.) and the Research School GROW (A.O., P.L., H.S.).

## Authors contributions

F.V. and H.S. designed and supervised the project and shared joint last authorship in the manuscript. A.O. and R.K. designed, performed most experiments, coordinated collaborations with other authors (P.L., M.G., H.P., M.M.), and A.O. and R.K. wrote as shared first joint authorship in the manuscript. P.L. designed, assigned, and supplied transcriptomic computational analysis. R.K., F.V. and H.S. determined PANTHER pathway gene-expression analysis on transcriptomic data. Further, Tfam knockdown experiments as quantitative and qualitative analysis, imaging embryos, oxygen consumption rate measurements were performed by A.O., R.K. in the manuscript. M.G. assisted on the design and scientific input on the manuscript. H.P., and M.M. assisted on the design and scientific input in the manuscript and were coordinating the housing and injections of the zebrafish embryos. All authors contributed ideas to the project.

## Supplementary information

Supplementary material is available for this article.

